# How Transcranial Magnetic Stimulation over Early Visual Cortex impacts short-term memory precision and guess rate

**DOI:** 10.1101/079178

**Authors:** Rosanne L. Rademaker, Vincent G. van de Ven, Frank Tong, Alexander T. Sack

## Abstract

Neuroimaging studies have demonstrated that activity patterns in early visual areas predict stimulus properties actively maintained in visual short-term memory. Yet, the mechanisms by which such information is represented remain largely unknown. In this study, observers remembered the orientations of 4 briefly presented gratings, one in each quadrant of the visual field. A 10Hz Transcranial Magnetic Stimulation (TMS) triplet was applied directly at stimulus offset, or midway through a 2-second delay, targeting early visual cortex corresponding retinotopically to a sample item in the lower hemifield. Memory for one of the four gratings was probed at random, and participants reported this orientation via method of adjustment. Replication errors were smaller when the visual field location targeted by TMS overlapped with that of the cued memory item, compared to errors for stimuli probed diagonally to TMS. This implied topographic storage of orientation information, and a memory-enhancing effect at the targeted location. Furthermore, early pulses impaired performance at all four locations, compared to late pulses. Next, response errors were fit empirically using a mixture model analysis to characterize memory precision and guess rates. Memory was more precise for items proximal to the pulse location, irrespective of pulse timing. Guesses were more probable with early TMS pulses, regardless of stimulus location. Thus, whereas TMS administered at the offset of the stimulus array might disrupt early-phase consolidation in a topographically unspecific manner, TMS also boosts the precise representation of an item at its targeted retinotopic location, perhaps by increasing attentional resources or by injecting a beneficial amount of noise.

## Introduction

Humans sense the world in a highly visual fashion – the flow of information from the eyes gives rise to an ostensibly effortless and seamless picture of our external environment. Despite its apparent simplicity, visual perception requires the brain to form an ongoing internal representation of all the information we are perceiving and perceived just moments ago, even if this information can no longer be sensed directly. Short-term memory takes center stage in the process of cognition by allowing relevant information to be kept online for further computation, serving as an indispensible buffer for human thought. Here, we investigated short-term memory for visual information and the role of early visual cortex during the maintenance of such information.

How might the brain meet the computational demands associated with the maintenance of information to which it no longer has access? The act of keeping visual memories online involves a network of frontal [1,2] and parietal [3–5] regions, as well as visual areas that were involved when the information was originally sensed [6–9]. The coordinated effort of higher-level and sensory brain regions during the short-term retention of visual information is believed to be flexible and goal dependent [10]. The dominant view in the literature on short-term memory is that higher-level areas recruit sensory areas that are specialized in processing the sensory analogs of specific mnemonic contents [11–14].

It has been suggested that sensory recruitment during visual memory is achieved in a spatially global and non-retinotopic manner: While people remembered an orientation presented in the left visual field, this orientation was decodable from patterns of functional Magnetic Resonance Imaging (fMRI) activity originating from *both* ipsi- and contralateral primary visual cortex (V1) [1,15]. However, the task used in these studies did not require subjects to maintain the relevant feature (here: orientation) bound to any specific location on the screen. Therefore, the lack of retinotopic recruitment could also be interpreted as a spread of feature-based attention [16–18]. Conversely, memory for visual information does depend on retinotopically specific representations when stimulus location is made relevant, and the explicit binding of stimulus contents to a particular location is required to perform a task. For example, location matters when people remember objects in a scene [19,20], when two orientations are presented one in each hemifield [8], or when location is made salient by a spatial transformation during memory [21,22].

To directly probe the causal role of sensory areas during the retention of visual stimuli, as well as the spatial extent of such recruitment, brain processes during memory can be actively altered by means of Transcranial Magnetic Stimulation (TMS). Previous work with TMS has provided support for both the necessity of visual sensory recruitment [23], as well as retinotopically specific maintenance of representations in these areas [24,25]. However, previous TMS studies demonstrating that the spatial extent of memory representations was confined in a retinotopic manner suffered some drawbacks: Specificity was only found very early during retention, probably during encoding [24], or was measured indirectly via the qualitative judgment of phosphenes [25]. Thus, while these studies suggest that brain stimulation has the potential to impact visual memories at the level of sensory representations, it remains to be seen whether retinotopically specific effects on performance can be found when TMS is applied outside of the range of sensory encoding.

While sensory recruitment during visual short-term memory has been well documented in studies measuring blood oxygenation with fMRI, the functional role of such recruitment is much less understood. In addition to the issue of retinotopic specificity, a second unanswered question concerns the functional role of sensory recruitment during short-term memory maintenance. Given that sensory areas can represent information with a degree of precision not easily achieved by less specialized areas, we hypothesized that their role during memory is to maintain high-precision representations.

Here, questions of *specificity* and *functional relevance* were addressed by applying TMS over occipital cortex while participants were remembering four oriented gratings, presented one in each quadrant of the visual field. By cuing memory based on spatial location, this task encouraged participants to encode and retain orientation information at the spatial locations at which they were presented. This design binds object identity to spatial position [26] allowing us to test for retinotopically specific recruitment of visual sensory cortex during short-term visual memory. To probe the functional role of sensory areas during the maintenance of orientation information, we employed a novel combination of methodologies: We combined TMS with rigorous psychophysical testing using the method of adjustment. This procedure involved collecting many trials per participant, calculating the angular deviation between the reported and true orientation on each trial, and analyzing the resultant error distributions by fitting a mixture model [27]. A mixture model characterizes memory errors as having two underlying sources: response variability and probability of uniform responses [27]. We applied this model to evaluate, respectively, the effects of TMS on memory precision and the effects of TMS on the likelihood of successful memory maintenance (i.e. “guess-rate”).

Pulses were applied at two different time intervals to check for potential differences between processes occurring at the tail end of encoding, and processes occurring well within the retention phase. Previous work showed that behavioral effects of visual cortex TMS during short-term memory depends on the timing of TMS [28]. In this study, participants remembered a circle with two lines extending from its center, forming a wedge. After a two-second delay participants judged whether a target dot appeared inside or outside of the remembered wedge. When TMS over visual cortex was applied at the end of the delay, responses were faster compared to vertex or no TMS stimulation. By contrast, TMS over visual cortex applied at the onset of the delay slowed response times [28]. Because brain stimulation interacts with ongoing neural activity at the stimulated region [29,30] we expected that TMS at different stages of retention (encoding and maintenance), would differentially impact behavioral outcomes.

Our primary hypothesis was that TMS over visual cortex would impact memory precision (but not guess-rate) in a retinotopically specific manner. A change in precision is expected based on the clear link that exists between information contents in visual cortex (as indexed by classification performance) and mnemonic resolution (as indexed by behavior), with more information predicting higher behavioral precision [6] [31]. One means by which TMS could impact behavior is by locally injecting random noise. Such noise could act to reduce the amount of information at the TMS location in visual cortex, and consequently negatively impact behavioral precision. Alternatively, at low levels of noise behavioral precision could improve: If weak neural signals are below (firing) threshold, a small amount of noise can push the intensity of these weak signals above threshold, enhancing signal discriminability – an idea known as ‘stochastic resonance’. Note that effects of noise on information transfer are non-linear, because with no (or too little) noise a threshold will not be reached, while too much noise will drown out the signal. Indeed, visual sensitivity is improved with low-intensity (below phosphene threshold) TMS stimulation [32], and low-intensity TMS facilitates behavior based on weak, but not strong, neural signals [33]. Mnemonic signals likely rely on weak sub-threshold signals, and low-intensity visual cortex TMS during short-term memory has indeed been shown to benefit behavior [25,28,34].

Here we found that TMS applied during the short-term retention of orientation stimuli improved overall behavioral performance in a retinotopically specific manner. This localized improvement mirrored by an increase in memory precision at the TMS location. Furthermore, TMS early during retention – at the tail end of encoding – resulted in a global, non-retinotopic, reduction in overall performance compared to TMS late during retention. This reduction in performance was mirrored by an increase in the likelihood of guess-like responses.

## Methods

### Participants

Eight participants were recruited from Maastricht University (5 females; mean age = 25.13 years, SE = 0.81). All had normal or corrected-to-normal vision, provided written informed consent, and passed a medical screening based on published safety guidelines [36] overseen by an independent medical supervisor. The medical ethics committee of the Maastricht University Medical Centre approved the study. With exception of one of the authors, participants received monetary reimbursement.

### Overall study design

We combined functional and anatomical MRI, with neuro-navigated TMS during a psychophysical short-term memory task (Fig 1A). Neuroimaging was utilized for the purpose of neuro-navigation [37], allowing constant TMS stimulation sites across multiple psychophysical sessions based on individually localized visual cortical activity. During the first (fMRI) session, anatomical and functional localizer data were obtained. The second (TMS) session determined the exact TMS target points and intensities to be used throughout subsequent experimental sessions, and participants furthermore performed 160 practice trials of the main psychophysical task.

**Fig 1.**
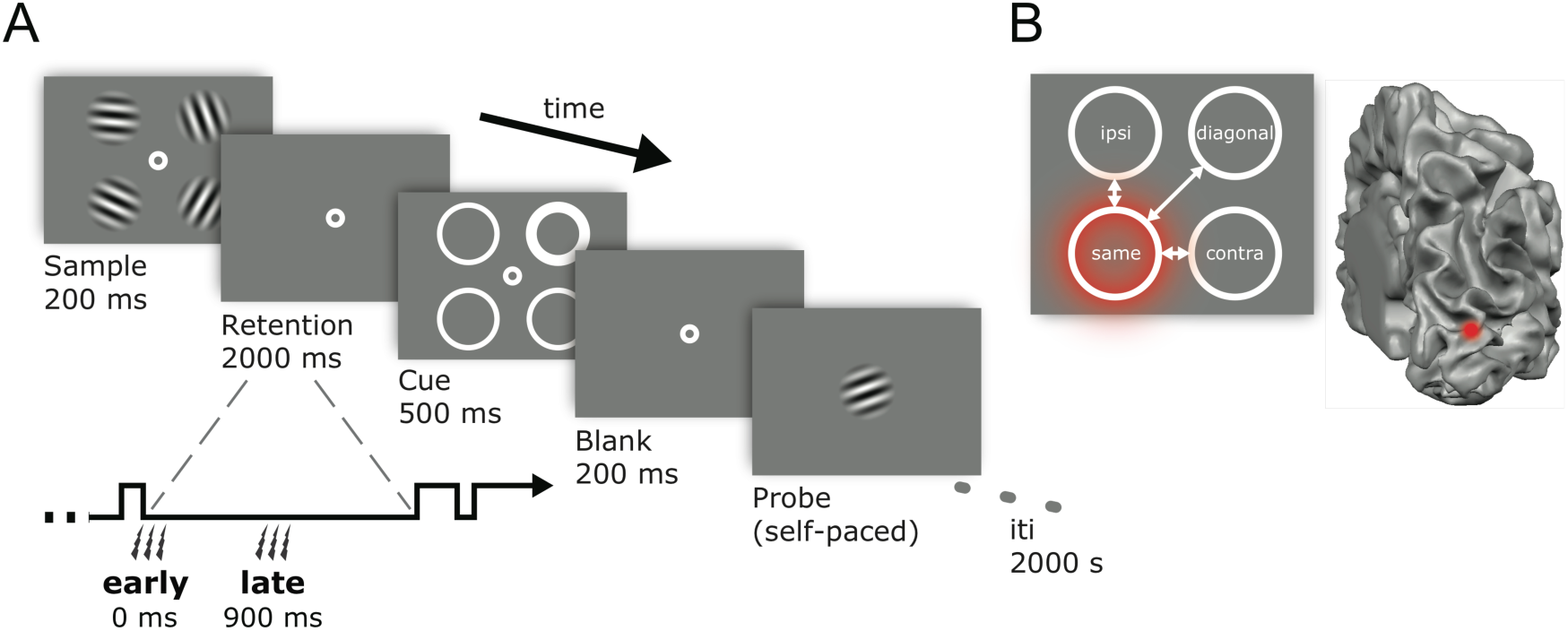
Trial sequence and relative locations. **(A)** Participants viewed a sample array displaying 4 randomly chosen orientations, and remembered these over a two-second delay. During this retention interval, participants received 3 pulses of real or sham TMS over their left or right hemisphere. The pulses arrived either directly at the offset of the sample array, or midway during the retention interval. A cue array indicated which of the four orientations was probed for recall, and after a short blank, participants rotated a test grating via button presses to match the cued orientation. **(B)** Responses at the four visual field locations were analyzed according to their position relative to the pulse. The visual field position targeted by the pulse could overlap with the memory item that was cued (‘same’), the cued item could be contralateral to the affected visual field location (‘contra’), it could be ipsilateral to it (‘ipsi’), or diagonal to it (‘diagonal’). In the example depicted here, the dorsal part of visual cortex in the right hemisphere is stimulated, targeting the lower-left visual field. Consequently, the upper left position becomes ‘ipsi’, the upper right position ‘diagonal’, the lower left ‘same’, and the lower right position ‘contra’ – relative to the visual field location affected by the TMS pulse.

During the last 5-6 sessions psychophysical data was collected while applying TMS over visual cortex. A TMS coil (real or sham) was placed over either the left or the right dorsal part of early visual cortex (V1/V2). Sham TMS was used to control for attentional biasing effects that can arise simply from “clicking” sounds at different points in time [37]. For half of our participants, a single session consisted of 4 blocks of 80 short-term memory trials per block, of which three blocks involved triple-pulse TMS stimulation at 10 Hz, and one block involved triple-pulse sham stimulation at 10 Hz. Target hemisphere (left or right) and type of stimulation (real or sham) were counterbalanced over blocks, sessions, and participants. The other half of participants underwent the same procedure, with one exception: they performed the blocks of sham-stimulation separately, several months after completing the real TMS sessions.

### MRI measurements

#### MRI acquisition

Scanning was performed at the Maastricht Brain Imaging Center (M-BIC) on a 3.0-Tesla Siemens MAGNETOM Allegra scanner using a standard birdcage head coil. A high-resolution 3D anatomical T1-weighted scan was acquired from each participant (FOV 256 × 256, 1 × 1 × 1 mm^3^ resolution, 192 slices, MPRAGE). To measure BOLD contrast, standard gradient-echo echoplanar T2*-weighted imaging was used to collect 28 slices covering the entire occipital lobe. Scan parameters for six participants were: TR, 2000 ms; TE, 30 ms; flip angle, 80º; FOV 192 × 192; slice thickness, 3 mm (no gap); in-plane resolution, 3 × 3 mm^2^. For two others scan parameters were: TR, 2000 ms; TE, 30 ms; flip angle, 90º; FOV 256 × 256; slice thickness, 2 mm (no gap); in-plane resolution, 2 × 2 mm^2^.

#### MRI data analysis

Preprocessing and analysis of the anatomical and functional MRI data were performed using BrainVoyager QX software (version 2.3.0.1750, Brain Innovation, Maastricht, the Netherlands). All anatomical data underwent inhomogeneity correction of signal intensity across space, and a tissue contrast enhancement using a sigma filter (7 cycles, range 5). Automatic grey-white matter segmentation was performed, after which manual corrections were made to improve segmentation over occipital cortex. The borders of the two resulting segmented sub-volumes were tessellated to produce surface reconstructions (folded meshes) – one for each hemisphere. These reconstructions were created to recover the exact spatial structure of the cortical sheet and to improve the visualization of anatomical gyrification.

#### functional MRI

Localizer stimuli in the scanner were generated using MATLAB 7.10.0 (R2010a) and the Psychophysics Toolbox [38]. Stimuli consisted of 5 Hz flickering black-and-white checkerboards (1º radius) presented 4º from fixation in either the lower left or lower right (randomly interleaved) quadrant of the screen against a uniform grey background (55.86 cd/m2). Stimulus locations encompassed the same visual field position as the two lower Gabor patches in the short-term memory task (Fig 1A). Stimuli were viewed through a mirror system on a back-projected screen (1024 × 768 resolution, 60 Hz refresh rate) at a distance of 66 cm in an otherwise darkened scanner room. We presented our localizer in two 5-minute functional runs, alternating twelve times between a 12 second fixation period and a 12 second stimulus period. Participants fixated throughout (0.5º white bull’s eye) monitoring occasional dimming of the checkerboard (~5 times per block), which was detected 43.18% of the time (SE=0.04).

After discarding the first 4 functional volumes, we applied automated 3D motion correction, slice-scan time correction (sinc), and high pass temporal filtering (using a GLM-Fourier basis set with 2 cycles) to correct for slow temporal drifts in signal intensity. No spatial or temporal smoothing was applied directly. Next, the fMRI data was aligned to the within-session anatomical scan via rigid-body transformations, with all automated alignment carefully inspected and manually fine-tuned when necessary. Functional data from both runs were combined and analyzed using a general linear model (GLM; [39]). Activity for the left and right hemispheres was based on the statistical contrast between BOLD signal [40] elicited by visual stimulation in the lower right versus lower left visual field respectively.

### Localization of the TMS target points

The anatomical reconstruction of a participant’s head was co-registered with the participant’s head in real space using stereotaxic data recorded with an ultrasound digitizer. Neuro-navigation was used to manually maneuver the TMS coil relative to a participant’s skull, while seeing in real-time their computer generated anatomical surface reconstruction with functional localizer activity superimposed. TMS target points were defined to lie within this region of activation. Specifically, each target point was chosen as posteriorly as possible within this region of activation, while still eliciting a phosphene overlapping the visual field location where stimuli would be presented (3–5º from fixation in lower left and right quadrants). Each TMS target point was indicated on the cortical surface reconstruction by a digital marker and saved to guide neuro-navigation for all future sessions. Phosphene thresholds were determined for the left and right hemispheres individually (at the saved TMS target points) and kept constant throughout all future sessions. Two participants did not experience phosphenes: target points were chosen at the peak-activity determined with fMRI, and stimulation intensity was set at the average intensity of other participants in the study.

### TMS protocol

Biphasic TMS pulses were delivered by means of a figure-of-eight coil (MCB70) and a MagPro R30 stimulator (Medtronic Functional Diagnostics A/S, Skovlunde, Denmark). This setup allows for pulse strengths (defined as the rate of change in the magnetic pulse) up to 148 A/µs at 100% of stimulator output, and 52 A/µs at 35% stimulator output. Pulses were applied at 80% of phosphene threshold to ensure that participants did not perceive visual stimulation (i.e. phosphenes) due to TMS.

Participants received 240 TMS pulses (3 pulses * 80 trials) during each run of the short-term memory task, and performed a total of 16 runs. The average pulse intensity used was 34.44% (SE=0.54) of maximum stimulator output, with no significant difference between the two hemispheres (mean left = 35 %, mean right = 33.88%, t = 1.386, p = 0.208).

### Short-term memory task

#### Stimuli

Experimental stimuli were generated with MATLAB 7.10.0 (R2010a) and the Psychophysics toolbox [38] under Windows XP and viewed in a dark room on a luminance-calibrated 19” Dell TFT monitor (1280 × 1024 resolution, 60Hz refresh rate). Communication between the experimental pc and stimulator was established using PortTalk V2.0 (Beyond Logic). Stimuli consisted of randomly oriented gratings with a spatial frequency of 2 cycles/º, a diameter of 2º, a 20% Michelson contrast with a wide Gaussian envelope (sd = 2º) and presented on a uniform grey background that shared the mean luminance of 40.23 cd/m2. Stimuli were presented at four fixed locations around a central fixation point at an eccentricity of 4º. Participants viewed the stimuli from 57 cm, and were instructed to maintain steady fixation throughout aided by a centrally presented white bull’s eye (0.5º diameter). A chinrest, forehead rest, and the tight placement of the TMS coil against the back of the head assisted in maintaining head stability.

#### Procedure

Observers were presented with a 200 ms sample array of 4 to-be-remembered gratings (Fig 1A). Each grating had an independently chosen random orientation (0-180º), with the only constraint that simultaneously presented orientations differed by >10º. During a 2-second retention interval a TMS triple-pulse was applied at 10 Hz (thus taking 200 ms) either directly following the offset of the sample array, or midway through the retention interval (the first pulse occurring 900 ms into the interval). Next, 4 spatial cues appeared for 500 ms outlining the locations of the previously presented sample stimuli. A 0.15º wide white circle probed the location of the grating to be reported from memory. Non-target locations had 0.05º outlines. After 200 ms a test grating was presented centrally at an initially random orientation. Participants used separate buttons on a keyboard to rotate the test grating clockwise or counterclockwise to match the probed orientation. Cue location was conceived of as counterbalanced (with each location probed an equal amount of times), but due to unforeseen circumstances the cue location was chosen randomly, without exactly equal amounts of trials at each location (see also supplementary information).

### Analyses

Due to anatomical constraints, TMS can only be applied over the dorsal (and not ventral) part of visual cortex. Thus, TMS can only target the two lower (and not upper) of the four stimulus-locations probed during our memory task. Previous studies have relied on this anatomical feature to contrast behavioral performance between a ‘TMS quadrant’ (usually the lower left) and a ‘control quadrant’ (usually the upper right)[24,41–43]. In the current task, the memory target could be cued in all four visual field quadrants. Consequently, data were analyzed according to the relationship between (1) the visual field location targeted by the TMS pulse, and (2) the visual field location probed for recall. To illustrate this analysis based on “relative location”, let’s assume that during a given block of trials the right hemisphere was stimulated with TMS, targeting the lower-left visual field (see also Fig 1B). If the target was subsequently cued in the same lower-left quadrant, both TMS and stimulus were presented at the “same” relative location. On trials where the target was cued in the upper-left quadrant (ipsilateral to the TMS-pulse location) the relative location was “ipsilateral”. Similarly, a target cued in the lower-right quadrant is at a “contralateral” relative location, whereas a target cued in the upper-right quadrant is at a “diagonal” relative location. The same logic can be applied when the coil is moved to the left hemisphere, targeting the lower-right visual quadrant.

Note that the *diagonal* location represents the *control quadrant*, as frequently employed in TMS studies of visual cortex. Such a control quadrant carries two major advantages over sham stimulation. One is that the diagonal location is probed randomly inteleaved with trials probing other locations, ensuring that participants are in the same general state during both trial types. Second is that real TMS is applied even when participants are probed at the control quadrant, ensuring that the acoustics and tactile experience during these trials perfectly matches that of other probed locations.

To separately estimate the precision of memory for successfully remembered items and the likelihood of memory failure we adopted a mixture-model approach following the work of Zhang and Luck (2008). This model summarizes data from method-of-adjustment tasks in a way that reflects the underlying assumptions of the model: on some trials items are remembered with a certain degree of precision, whereas on other trials items are not available for recall resulting in random guesses. This idea was implemented by fitting a circular Gaussian-shaped model to the distribution of orientation errors (actual orientation minus reported orientation) for each condition of interest. The model consisted of two key parameters: One is the standard deviation (*SD*), or width of the circular portion of the distribution, assumed to reflect the precision of short-term memory for successfully remembered items (with better precision indicated by a smaller *SD*). Two is the relative proportion of area under the curve corresponding to a uniform distribution (*p-Uniform*), which captures the extent to which the entire distribution needed to be translated along the y-axis to account for the frequency of guess-like responses, and is assumed to reflect the probability of guessing responses. We rely on these summary statistics throughout this paper because they provide a useful way to capture broad trends in the data and because they may signify distinct types of errors. However, it is important to acknowledge that the mapping between these summary statistics and underlying sources of error in the short-term memory system rely on assumptions regarding the exact nature of short-term memory performance, and that competing models have been proposed (e.g., [44–48]).

## Results

### Absolute performance

Experimental manipulations consisted of (1) probing memory for items at various relative distances from the visual field location targeted by TMS, and (2) applying TMS early or late during memory retention. Moreover, TMS was applied in a counterbalanced fashion either over the left- or right hemisphere, which was not expected to impact performance. By performing a 3-way within-subjects ANOVA (4 relative locations × 2 pulse timings × 2 stimulated hemispheres) on the absolute errors (absolute difference between reported and true orientation) we confirmed that the stimulated hemisphere (left or right) did not affect memory accuracy (*F*_(1,7)_ < 0.001; *p* = 0.986).

Proximity of a memory item to the visual field location targeted by TMS while items were being maintained in memory had a facilitative effect on memory performance, as indexed by smaller errors for items proximal to the pulsed location (main effect of relative location; *F*_(3,21)_ = 3.951; *p* = 0.022; Fig 2). Six post-hoc ANOVA’s were performed to investigate the origins of this main effect of location, each comparing performance at two locations (each 2 relative locations × 2 pulse timings × 2 stimulated hemispheres). The location effect depended on the difference between trials on which the TMS pulses and probed location overlapped (‘same’ condition) versus when they were furthest apart (‘diagonal’ condition, i.e. the control quadrant) (*F*_(1,7)_ = 5.598; *p* = 0.050).

**Fig 2.**
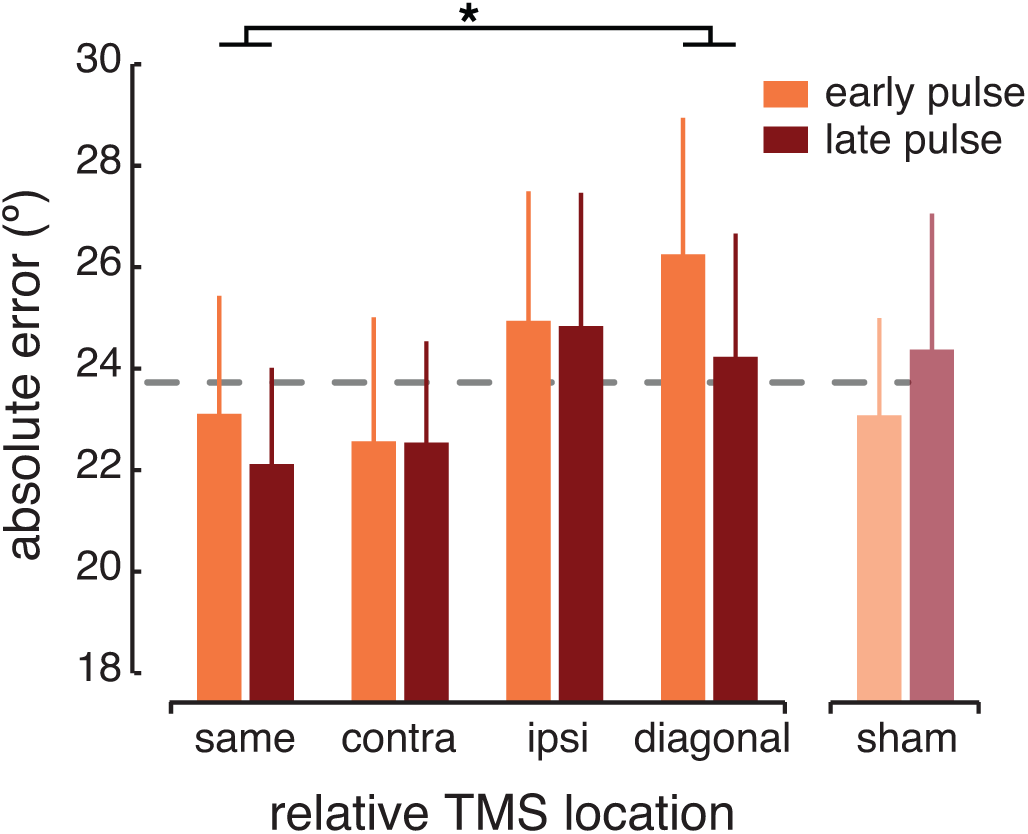
Absolute performance relative the TMS location. When a memory item was cued at the same location as targeted by the TMS pulses, the absolute response error was smaller than when a memory item was probed diagonally to the pulses. This indicated better performance on trials where memory cue location and TMS location overlap (compared to a cue diagonal to TMS). Early pulses resulted in worse performance compared to late pulses, irrespective of the cued visual field location. Trails during which TMS was administered are shown here after collapsing across both hemispheres, since stimulation site (left or right hemisphere) did not affect participant’s performance. Sham data is shown collapsed across all conditions (dashed grey line), and separately for early and late sham pulses (orange and red bars). The time point at which audible sham clicks were delivered did not impact behavior. When data were collapsed across both hemispheres (as depicted here), a 4x2 ANOVA (relative location by pulse timing) revealed statistically reliable performance differences between the four visual field locations (*F*_(3,21)_ = 3.483; *p* = 0.034) and a marginally significant effect of pulse timing (*F*_(1,7)_ = 4.820; *p* = 0.064). Error bars depict + 1 SEM.

Comparing the two pulse-timings showed that applying TMS pulses early, directly at the offset of the stimulus display, resulted in larger errors than TMS applied midway through the retention interval (main effect of pulse timing; *F*_(1,7)_ = 6.359; *p* = 0.040).

Audible ‘clicks’ from a TMS machine can have differential attentional cuing effects when presented at different time points [37], with the potential to influence behavior differently early and late during retention. Because sham TMS has no neural effects, we collected a total of 320 sham trials (160 per pulse timing) to investigate potential effects of timing. While this is the right amount of trials for looking at timing effects, note that this is much less than the 1280 trials collected during real TMS, precluding analyses of the sham data in a manner truly identical to analyses of the real TMS data. We analyzed sham data by first looking at our condition of interest, pulse timing, while collapsing across all other factors (location and hemisphere). The time point at which sham pulses were applied did not impact performance (*t*_(7)_ = 0.647; *p* = 0.538). We also compared performance for absolute location (comparing upper left, upper right, lower left, and lower right locations) and hemisphere (left and right) collapsing across the remaining factors. Again, no differences were observed (*F*_(3,21)_ = 0.459; *p* = 0.714 and *t*_(7)_ = 1.013; *p* = 0.345, for location and hemisphere respectively). The finding that there were no differences between any of the conditions on sham trials is important, since our manipulations could have had unintended differential attentional-cuing effects, mimicking the neural effects probed via TMS.

### Mixture-model results

To gain a deeper understanding into the functional role of early visual cortex involvement during the maintenance of visual memories, we fit a mixture model to data from each combination of location and pulse timing conditions (collapsed across hemispheres). This allowed us to decompose the absolute errors into (1) memory precision as indexed by the mixture model *SD* (with a smaller *SD* indicating better precision), and (2) the likelihood of guess-like responses indexed by *p-Uniform* (with a larger *p-Uniform* indicating more guessing) [27].

When a memory target was probed at a location proximal to the location targeted by TMS, memory recall was more precise (4 relative locations × 2 pulse timings ANOVA, main effect of relative location; *F*_(3,21)_ = 4.102; *p* = 0.019; Fig 3A). Specifically, six post-hoc ANOVA’s (comparing all possible pairs of relative locations) showed that memory was more precise on trials where the pulses and probed memory item overlapped, i.e. ‘same’ condition, (*F*_(1,7)_ = 7.974; *p* = 0.026) or were ‘ipsilateral’ to one another (*F*_(1,7)_ = 7.658; *p* = 0.028), compared to trials on which the pulses and probed item were furthest apart (‘diagonal’, control quadrant).

**Fig 3.**
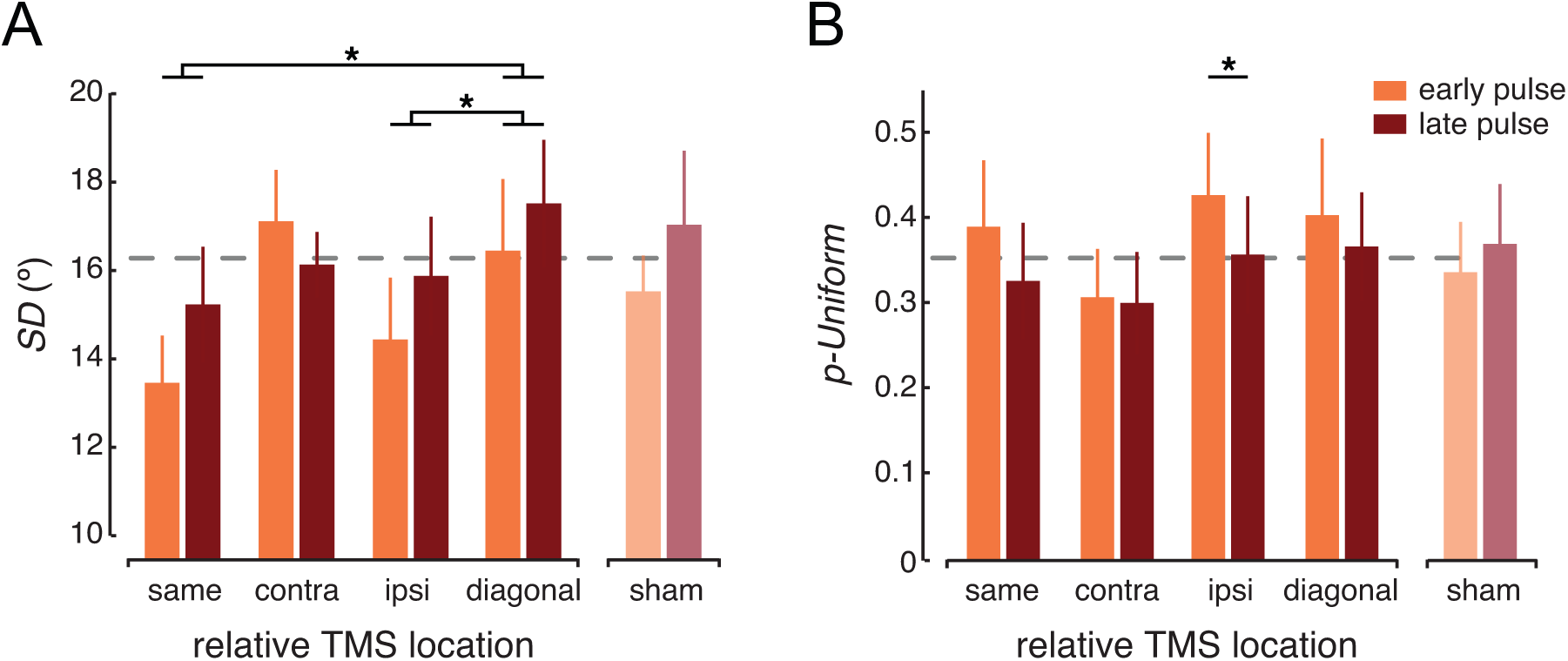
Model fits of TMS data. **(A)** Memory precision is represented by the mixed-model *SD*, with a smaller *SD* indicating more precise memory. Memory was most precise when the location of a cued memory item overlapped with the location at which TMS was applied, or was ipsilateral to it, compared to diagonally to pulses. **(B)** When pulses were delivered early during the retention interval, participants were more likely to guess compared to pulses delivered late, irrespective of their retinotopic location relative to TMS. Parameter estimates were obtained by finding the bestfitting mixture-model (centered on 0º error of report, based on the mixed model analysis) for the frequency distribution of each condition, using a bin width of 12º (mean R2 = 0.894 + 0.029). Bin size was chosen to maximize the mean R2 values across the different experimental conditions. Data were collapsed across hemispheres (left and right stimulation) before fitting in order to achieve a large enough number of trials per condition to obtain reliable fits. Error bars depict + 1 SEM.

No retinotopic specificity was found for the probability of uniform responses, which did not differ significantly between the four visual field locations (*F*_(1,7)_ = 1.831; *p* = 0.172). However, participants were more likely to show guess-like responses when pulses were presented early during the retention interval compared to pulses presented midway through retention (main effect of pulse timing; *F*_(1,7)_ = 6.594; *p* = 0.037; Fig 3B). This increase in random responses occurred irrespective of the location at which the memory item was probed (no interaction; *F*_(3,21)_ = 0.712; *p* = 0.555).

Finally, sham data were fit with a mixture model for each condition of potential interest (pulse timing, and absolute location) separately, collapsing across all other conditions. Timing of the sham pulses did not affect memory variability (paired-samples *t*_(7)_ = 0.822; *p* = 0.438) nor the probability of guess-like responses (paired-samples *t*_(7)_ = 1.262; *p* = 0.247). We found no differences between the four absolute visual field locations for either memory precision (*F*_(3,21)_ = 0.976; *p* = 0.423) nor the probability of uniform responses (*F*_(3,21)_ = 1.272; *p* = 0.310).

### Memory performance across the visual field

A strength of our experimental setup was that the TMS coil was fixed over the skull, which, in combination with fMRI guided neuro-navigation, ensured that the targeted brain site remained stable relative to the four patches of retinotopic cortex excited by our stimuli. However, this setup had the obvious side effect that the stimuli were anchored onto four static visual field locations across all experimental trials. If perception and memory at these static visual field locations were anisotropic, this would complicate interpretation of our results. From the literature on basic human vision it is known that people are generally better at performing a visual task on stimuli in the lower, compared to the upper visual field. Luckily, such anisotropies are generally found for stimuli presented along the cardinal meridians, and less prevalent (or even absent) for stimuli presented at the obliques [49,50]. Nevertheless, any residual anisotropy would impact our results by biasing performance in favor of the lower half of the visual field, incidentally, the same half of the visual field targeted with TMS.

One crucial check was to investigate how memory on sham trials varied across the stimulus locations in the upper left, upper right, lower left, and lower right parts of the visual field. These results were already discussed above: During sham neither the absolute error (*F*_(3,21)_ = 0.459; p = 0.714), memory precision (*F*_(3,21)_ = 0.976; *p* = 0.423), nor the probability of uniform responses (*F*_(3,21)_ = 1.272; *p* = 0.310) varied as a function of visual field location probed. While this suggests that visual field anisotropies cannot explain our main findings, a null effect (absence of evidence) does not provide conclusive proof (evidence of absence). A second observation arguing against anisotropies is that, while results from the absolute performance seem to roughly align with the idea of upper-lower visual field anisotropies (Fig 2), results from the fitted memory precision (Fig 3A) imply no such distinction.

Finally, we also directly probed whether real TMS and sham TMS interact, as such an interaction would provide the most convincing evidence against the idea that upper versus lower visual field anisotropies (rather than TMS effects) might be driving performance differences at the four locations. In order to perform this direct comparison, we analyzed the absolute performance during sham in a manner identical to that of real TMS: performance was determined at each of the four visual field locations relative to the location ‘targeted’ by sham (for example, a sham coil over the right hemisphere ‘targeted’ the lower left visual field). Data for both sham and real TMS were collapsed across stimulated hemisphere, resulting in 160 and 40 observations per condition on average for real and sham TMS respectively. An ANOVA comparing the 2 TMS conditions (real and sham), 2 pulse timings (early and late), and 4 locations (‘same’, ‘contra’, ‘ipsi’, and ‘diagonal’) revealed a significant interaction between pulse timing and location (*F*_(3,21)_ = 3.752; *p* = 0.027). Note that neither TMS condition and location (*F*_(3,21)_ = 0.204; *p* = 0.893) nor TMS condition and timing (*F*_(1,7)_ = 2.007; *p* = 0.2) interacted significantly.

**Fig 4.**
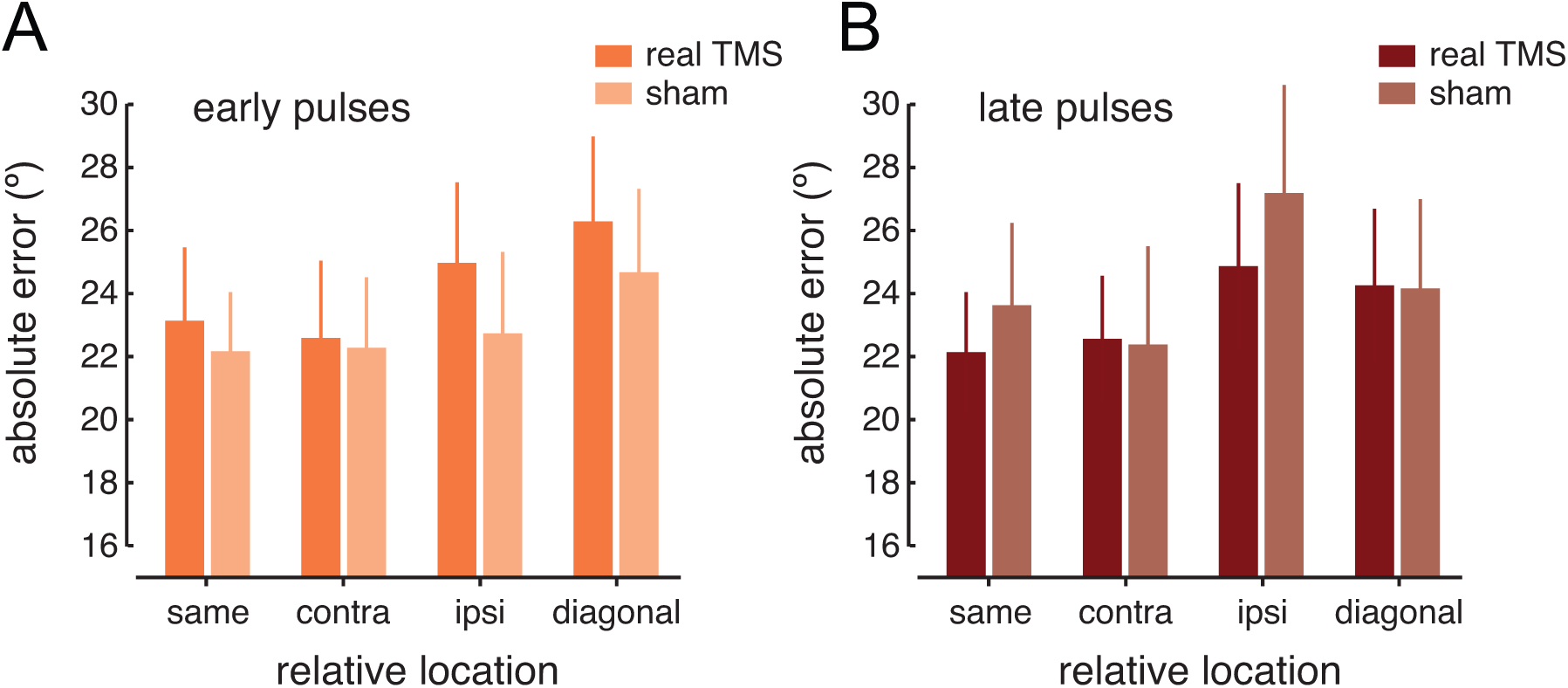
Absolute performance during real and sham TMS. **(A)** Performance when real or sham pulses were applied early during retention, directly at stimulus offset. **(B)** Performance for pulses (real or sham) applied midway through the retention interval. Error bars depict + 1 SEM.

Thus, direct comparison of real and sham TMS did not yield entirely conclusive results. One obvious problem is that sham data could have been more variable due to the smaller number of trials, making comparisons less robust (but also note that 40 trials per condition is still considerable when not performing any fitting procedures). A more pertinent problem is that performance during TMS is not expected to differ from performance during sham outside of the TMS-targeted location. In fact, when we only considered visual field locations where real TMS is expected to yield an effect (i.e. overlapping with, or ipsilateral to, the targeted location) the interaction between TMS condition and timing did reach significance (*F*_(1,7)_ = 5.975; *p* = 0.044), suggesting that real TMS and sham differentially affect performance at different time points of memory maintenance.

## Discussion

While participants were remembering four orientations, 10Hz triple-pulse TMS was applied over early visual cortex retinotopically corresponding to the location of one of the to-be-remembered items. Orientation recall differed between the four locations at which stimuli had been presented, with better performance at the location targeted by TMS compared to the location diagonal to TMS. Additionally, memory was impaired for early (directly at stimulus offset) compared to late (midway through retention) pulses. Replication errors were fit with a mixture model to reveal relative contributions of changes in memory variability on the one hand, and the probability of guessing responses on the other: Spatially specific improvements proximal to the pulse were attributed to reduced response variability, implying that memory precision can be improved locally by means of TMS. Global impairments for early compared to late pulses were due to an increased likelihood of guess-like responses, implying retinotopically aspecific disturbances due to TMS at the tail end of encoding. A cartoon-summary of these findings is shown in Figure 5. None of these findings were observed with sham TMS. While it is possible that performance at the four stimulus locations differed because of visual field anisotropies, such anisotropic contributions were likely not driving the observed TMS effects.

**Fig 5.**
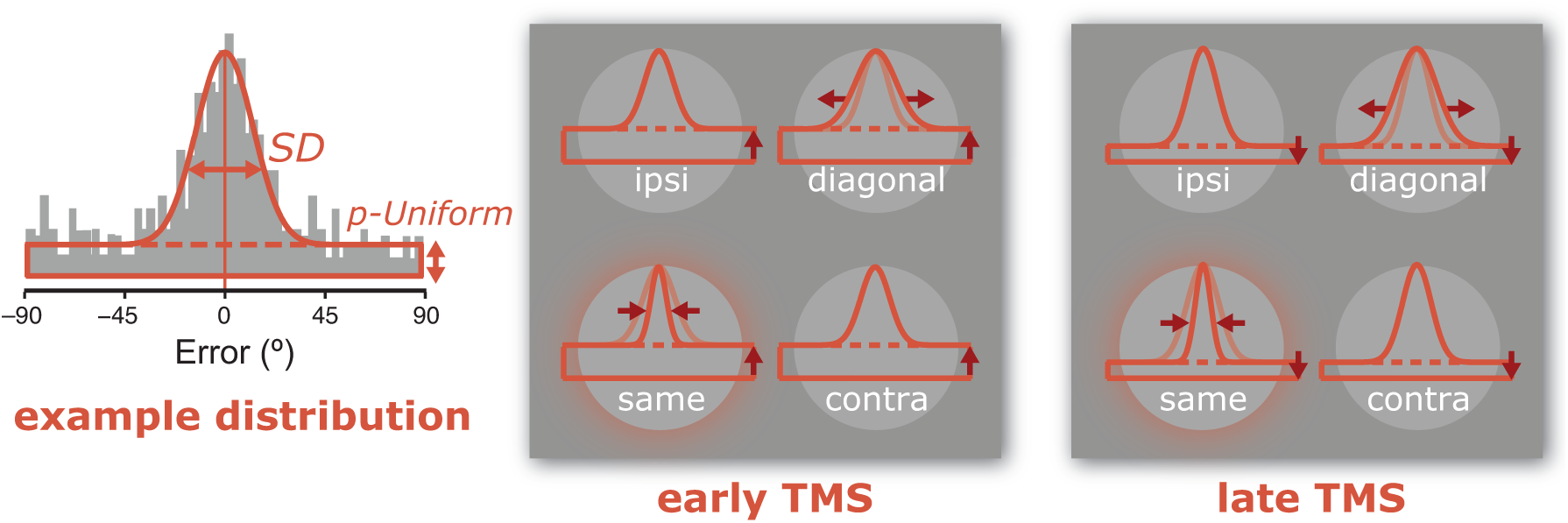
Cartoon summary of results. The left most panel depicts an example response distribution. The frequency of response errors (i.e. difference between cued orientation and response) is indicated by the height of the grey bars. Most responses are centered on a mean replication error of 0º, and the precision with which this replication is achieved is captured by the mixture model *SD* parameter (smaller *SD* indicates more precision). Some responses appear to be are random guesses, indicated by the probability of guesses or ‘uniform responses’ (< *p-Uniform* means less guessing). A mixture model fit to the response errors is depicted in thick orange lines. The two rightmost panels provide a cartoon summary of our main results (exaggerated and simplified for illustrative purposes). First, the main effect of relative location is shown by increased memory precision (*SD*) at the location targeted by TMS (‘same’) compared to the control quadrant (‘diagonal’) for both early and late TMS pulses (compare *SD* at four visual field locations). Second, the main effect of pulse timing is shown by increased guessing when pulses were applied early compared to pulses applied late irrespective of visual field location (compare *p-Uniform* between middle and right most panels).

Our task encouraged binding the memorized features to their visual field positions by making memory retrieval contingent upon spatial location. Limits on the spatial extent of sensory recruitment have been a matter of some debate, found in some cases [8] but not others [1,15]. Our findings of locally improved memory performance and precision provide support for retinotopically specific sensory recruitment during visual memory. What is more, this local improvement was present irrespective of the time point during which TMS was applied, demonstrating that TMS can impact visual short-term memories beyond the sensory encoding stage.

Why might early pulses lead to more guess-like responses than late pulses? Simple modulations of attention or distractibility due to the timing of the three auditory clicks emitted by the TMS coil could not explain this – using the same timing and sounds revealed no costs for early compared to late sham TMS. Guessing responses can result from forgetting, lapses of attention, or encoding failures. Since our early pulses were presented at the tail end of encoding [51–54], it is possible that TMS increased random guesses by prohibiting adequate encoding of the four stimuli. This finding is in line with previous work showing disruptions to visual memory when TMS was applied over visual cortex during the early stages of retention [24,28]. However, this earlier work observed retinotopically specific disruptions. Why might we find that early pulses affected performance across the visual field?

First, while stimulus orientations were chosen randomly and independently, accidental yet strong ensemble effects probably occurred on a portion of trials by virtue of our design. Anticipating the fixed trial-by-trial spatial location of memory stimuli, participants could have adapted a strategy relying on the constellation of the four orientations (as radial, concentric, isotropic, etc.) rather than storing features in a truly independent manner. Perceptual grouping of elements allows more of them to be stored in memory [55], and taking perceptual grouping and higher-order structures between items into account helps explain memory performance [56,57]. Thus, early TMS pulses applied while participants were extracting global shape-like representations could disrupt encoding of the whole ‘object’ through the disturbance of localized ‘object features’ (i.e. orientations), resulting in increased guesses for all orientations during retrieval. Such an ‘object-based’ short-term memory strategy might be achieved by convergent feed-forward and feedback processes at multiple stages of the visual hierarchy [58,59] – an intriguing possibility that could be tested empirically in the future.

Second, a higher likelihood of guessing responses for early compared to late pulses implied that different processes (or global cognitive ‘states’) were occurring at different stages of the retention interval. A brain ‘state’ can refer to many things like general arousal level, attention or inattention, being trained or untrained, adapted or unadapted, etc. Acknowledging that TMS interacts with the initial brain state helps frame prior studies applying TMS over visual cortex during short-term memory maintenance, unveiling both performance *decrements* [23,24,28], as well as *improvements* [28,34]. Specifically, it has been proposed that TMS may preferentially activate neurons in low initial activation states (i.e. low firing) relative to more active populations [30,60,61]. Alternatively, the effect of TMS on neuronal firing is monotonic, but behavioral effects (facilitation versus impairment) depend on non-linearity in the input response function of the sensory neurons [62]. Either way, additional mechanisms beyond global brain states must be assumed to account for *local* improvements in memory precision that exist independent (and in addition to) the *global* TMS timing effects reported here.

First, TMS might enhance processing of a memorized orientation *locally*. For example, we used low-intensity TMS, which could protect local populations of neurons at the TMS location against temporal decay by pushing weak signals above threshold. A related idea is that TMS enhancement depends on non-monotonic intensity responses [62] as mentioned above: The first basic premise is that TMS acts via a wholesale multiplication of neural responses. The second premise is that while remembering an orientation, the memory trace of that orientation in a population of orientation selective neurons is weak, with firing rates only slightly elevated above baseline. In terms of intensity response, the remembered orientation has a response that is slightly larger than that of not-remembered orientations. However, because of the nonlinearity of intensive response profiles, a wholesale multiplication by any factor due to TMS would result in higher signal-to-noise for the remembered orientation. Conceivably, such signal-to-noise benefits could be induced with TMS on a directly perceived orientation as well, as long as contrast remains lower that the inflection point of the contrast response function, and intensive responses undergo expansive nonlinearity.

Second, participants in our experiment simultaneously remembered multiple stimuli presented at *distributed locations*, each stimulus competing for processing resources. This raises the possibility that mnemonic representations interacted and that TMS yoked competition spatially, in favor of the targeted retinotopic location. By enhancing neural firing at one of multiple task-relevant locations, TMS might mimic spatial attention: It’s been shown that in the absence of a visual stimulus, spontaneous firing rates in V2 ad V4 were elevated when attention was directed at a location that fell within a cell’s receptive field [63]. Thus, TMS might act as a bottom-up implementation of an otherwise top-down biasing signal, potentially by means of a spatial gating mechanism that favors the boosted location during subsequent processing stages [64].

Note that the results reported here seem to indicate a mixture of both improved and impaired performance more proximal and distal to TMS respectively. Thus, TMS might be attenuating regulatory processes that mediate competition between orientation representations at distributed visual field locations. This can be framed in terms of a recent population-coding model of short-term memory [14,65,66] that assumes stimulus features (like orientation) are stored by probabilistic spiking activity in tuned populations of neurons. Critically, this model includes a broad normalization component by keeping the sum of the firing rates remains constant (across changes in set size or attentional prioritization, for example) [46,65]. Normalization describes neuronal responses as consisting of an input or ‘drive’, divided by the summed activity of a normalization pool [67,68]. The assumption that normalization occurs between all items held in memory could explain our findings via a multiplicative increase of the input drive of neurons at the TMS site, biasing the overall population activity in favor of the TMS location (and against non-TMS locations).

Alternatively, inhibitory interactions might exist between the four simultaneously remembered orientations: At a local level, TMS can release orientation representations from inhibitory interactions during a tilt illusion paradigm [69]. Likewise, in our study TMS could have acted to depress horizontal connections between spatially distributed representations, attenuating interference at the targeted location. In the brain, lateral connections between neurons in posterior visual areas with relatively large aggregate receptive field sizes [70], monosynaptic trans-colossal connections between the primary visual hemispheres [71], or top-down influences from for example prefrontal cortex [65,72] are all routes through which interactions between multiple and spatially distributed stimulus representations might arise.

These ‘local’ and ‘distributed’ hypotheses about improved processing at the TMS location, while thought-provoking, should be tested empirically in future work to ascertain their true value. To that, we’d like to add some additional considerations regarding the work presented here. First, noisy performance from a couple of participants precluded reliable fitting using Maximum Likelihood Estimation. Instead we binned the data in bins of 12º before deriving parameter estimates. While rather coarse, this bin size was chosen to maximize R2 values, and results were comparable for analyses using smaller bins (i.e. 8º or 10º). Error fitting performed in this way is purely empirical, unlike model fitting, and could even be considered the better choice here. However, it should be noted that when applying a maximum likelihood approach the directionality of effects was preserved, but neither precision differences across the visual field (*F*_(3,21)_= 1.837; *p* = 0.171), nor the pulse timing effects on guessing responses (*F*_(1,7)_ = 2.854; *p* = 0.135) remained significant.

Second, TMS effects are usually small, and people’s response to TMS highly variable, which is why TMS results generally benefit from large sample sizes. Instead, here we opted for an in depth psychophysical approach, spanning many sessions and trials. The strength of our design is that it allowed us to investigate the mechanistic underpinnings memory maintenance in visual cortex. Our findings dovetail previous reports of retinotopically [24] and temporally [28] specific effects of visual cortical TMS on short-term memory. In addition, our findings suggest that memory precision is affected retinotopically, while effects of pulse timing are likely a matter of global state-like processes. The trade-off however, was a relatively smaller number of participants.

Despite these cautionary notes, our results provide several consistent and intriguing findings. Combining the mixture-model with TMS offered novel insights into the role of early visual cortex during the short-term retention of visual items in memory, while affirming the involvement of these areas. We were able to uncover a double dissociation, showing local changes in memory precision at the TMS location, and global changes in guess rates contingent on pulse timing, with more frequent guesses for TMS applied at the tail end of encoding compared to midway during the delay. Our work adds to an existing literature demonstrating retinotopically specific sensory recruitment [8] and early visual cortex TMS impacting visual short-term memory [24,25,28].

## Acknowledgements

We would like to thank Sam Ling for valuable input during the analysis stage and long Skype sessions regarding the contents in the Discussion.

